# Temporally dissociable mechanisms of spatial, feature, and motor selection during working memory-guided behavior

**DOI:** 10.1101/2022.05.18.492385

**Authors:** Edward Ester, Rachel Weese

## Abstract

Working Memory (WM) is a capacity- and duration-limited system that forms a temporal bridge between fleeting sensory phenomena and possible actions. But how are the contents of WM used to guide behavior? A recent high-profile study reported evidence for simultaneous access to WM content and linked motor plans during WM-guided behavior, challenging serial models where task-relevant WM content is first selected and then mapped on to a task-relevant motor response. However, the task used in that study was not optimized to distinguish the selection of spatial versus non-spatial visual information stored in memory, nor to distinguish whether or how the chronometry of selecting non-spatial visual information stored in memory might differ from the selection of linked motor plans. Here, we revisited the chronometry of spatial, feature, and motor selection during WM-guided behavior using a task optimized to disentangle these processes. Concurrent EEG and eye position recordings revealed clear evidence for temporally dissociable spatial, feature, and motor selection mechanisms during this task, partially replicating yet also extending previous findings. More generally, our data reveal the existence of multiple WM selection mechanisms that belie conceptualizations of WM-guided behavior based on purely serial or parallel visuomotor processing.

---

A fundamental purpose of any memory system is to inform future decisions and actions. This is particularly important in the case of working memory (WM), a duration- and capacity-limited system that forms a temporal bridge between fleeting sensory phenomena and possible actions. Recent theoretical conceptualizations of WM have begun to emphasize the action-oriented nature of this system (e.g., Heuer et al., 2020; Olivers & Rolfsema, 2020; van Ede & Nobre, 2023), and recent empirical findings suggest that behavioral (Gonzalez-Garcia et al., 2020; Ohl & Rolfs, 2020), circuit-level (Pho et al., 2018), and systems-level (Chatham et al., 2014; van Ede et al., 2019a; Galero-Salas et al., 2021; Rak-Lubashevsky & Frank, 2021; Boettcher et al., 2021) mechanisms of WM storage and action planning are tightly interwoven.

Recent studies suggest that human observers can store multiple stimulus-response mappings in WM (see van Ede & Nobre, 2023 for a review), and that behaviorally relevant WM content can be selected in parallel with required actions. In one high-profile example, van Ede and colleagues (2019a) required participants to remember the orientations of two colored bars over a short delay. A color cue presented at the end of the delay informed participants which item would be probed for report, while the vertical tilt of the color-cue-matching bar informed participants which hand should be used for orientation recall (with clockwise and anti-clockwise tilted bars required right- and left-hand responses, respectively). Cleverly, van Ede and colleagues independently manipulated the physical location of the probed bar (i.e., left vs. right visual hemifield) and the tilt of the probed bar (i.e., requiring a left vs. right hand response), which allowed them to distinguish the selection of visual and motor information via lateralized signals measured in contemporaneous EEG recordings. Surprisingly, EEG signals associated with the selection of visual and motor information had nearly identical time-courses, suggesting that participants were able to select the task-relevant WM content and the appropriate response in parallel.

On the one hand, the findings reported by van Ede et al. (2019a) seem to challenge classic conceptualizations of human memory performance based on sequential processing stages where relevant information is first selected and then mapped onto appropriate outputs (e.g., Donders 1868/1969; Sternberg, 1969; Meyer et al., 1988). On the other hand, two aspects of the study performed by van Ede and colleagues (2019a) undermine this challenge. First, the electrophysiological signal that these authors used to track the selection of visual information stored in WM – lateralized alpha-band activity – is known to index covert spatial attention (e.g., Klimesch, 2012). Although van Ede and colleagues presented to-be-remembered stimuli in opposite visual hemifields (i.e., one bar appeared in the left visual field and another in the right visual field), participants were always probed to report a specific bar via a color cue and by adjusting a probe stimulus presented at fixation. Thus, participants could solve the task by storing color-orientation bindings independent of where these bindings appeared. While there is growing evidence that human observers automatically store spatial information in WM and use spatial information to select behaviorally relevant WM content (e.g., Foster et al., 2017; Groen et al., 2022), this leaves open the question of how non-spatial features are selected alongside motor plans during WM-guided behavior.

Second, and more importantly, the design used by van Ede and colleagues (2019a) confounded the selection of motor information with the selection of task-relevant feature information. Specifically, the vertical tilt of the retrospectively cued bar (i.e., clockwise vs. anticlockwise from vertical) instructed participants which hand should be used for recall (right or left, respectively). Thus, changes in lateralized beta-band EEG activity that these authors ascribed to motor selection could instead reflect feature selection (i.e., the clockwise or anticlockwise tilt of the to-be-recalled bar). Likewise, van Ede et al. (2019a) yoked response hand with probe rotation direction during recall. That is, right hand responses always produced clockwise rotations of the probe bar while left hand responses always produced anticlockwise rotations of the probe bar. Thus, the lateralized beta-band signals that these authors ascribed to motor planning could instead reflect a recall strategy based on feature information (e.g., “the probe stimulus must be rotated clockwise by 30°”) rather than motor planning per se. These factors, coupled with studies linking EEG beta synchrony with content-specific WM storage (summarized in Spitzer & Haegens, 2017) and studies linking beta local field potentials in non-human primates with top-down control over the contents of WM (summarized in Miller et al., 2018) motivate a fresh look at the role of beta oscillations in the selection of remembered feature and/or motor information.

In this study we sought further clarity on the chronometry of spatial, feature, and motor selection during WM-guided behavior. We recorded EEG and eye position data while human volunteers performed a retrospectively cued orientation recall task (see Figure 1 for a schematic). Following earlier work (van Ede et al., 2019a), we manipulated the physical location of the retrospectively cued orientation (i.e., left vs. right visual field) independently of response hand. Unlike prior work, we decoupled the selection of spatial, feature, and motor selection by (a) using a color cue to inform participants which item stored in WM was task-relevant and which hand should be used for recall, and (b) delaying orientation recall until one second after cue onset. To preview our findings, spectral analyses of EEG data and concurrently recorded eye position data time-locked to retrocue onset revealed evidence for the simultaneous selection of spatial and motor information following the onset of an informative retrocue, replicating van Ede and colleagues (2019a). Critically, above-chance decoding of stimulus orientation was delayed until the onset of the response display, demonstrating that feature selection is temporally decoupled from spatial and motor selection. These findings challenge models of WM-guided behavior based on purely serial or parallel selection of visual and motor information, and instead point to the existence of multiple temporally distinct selection mechanisms.

**Figure 1.**
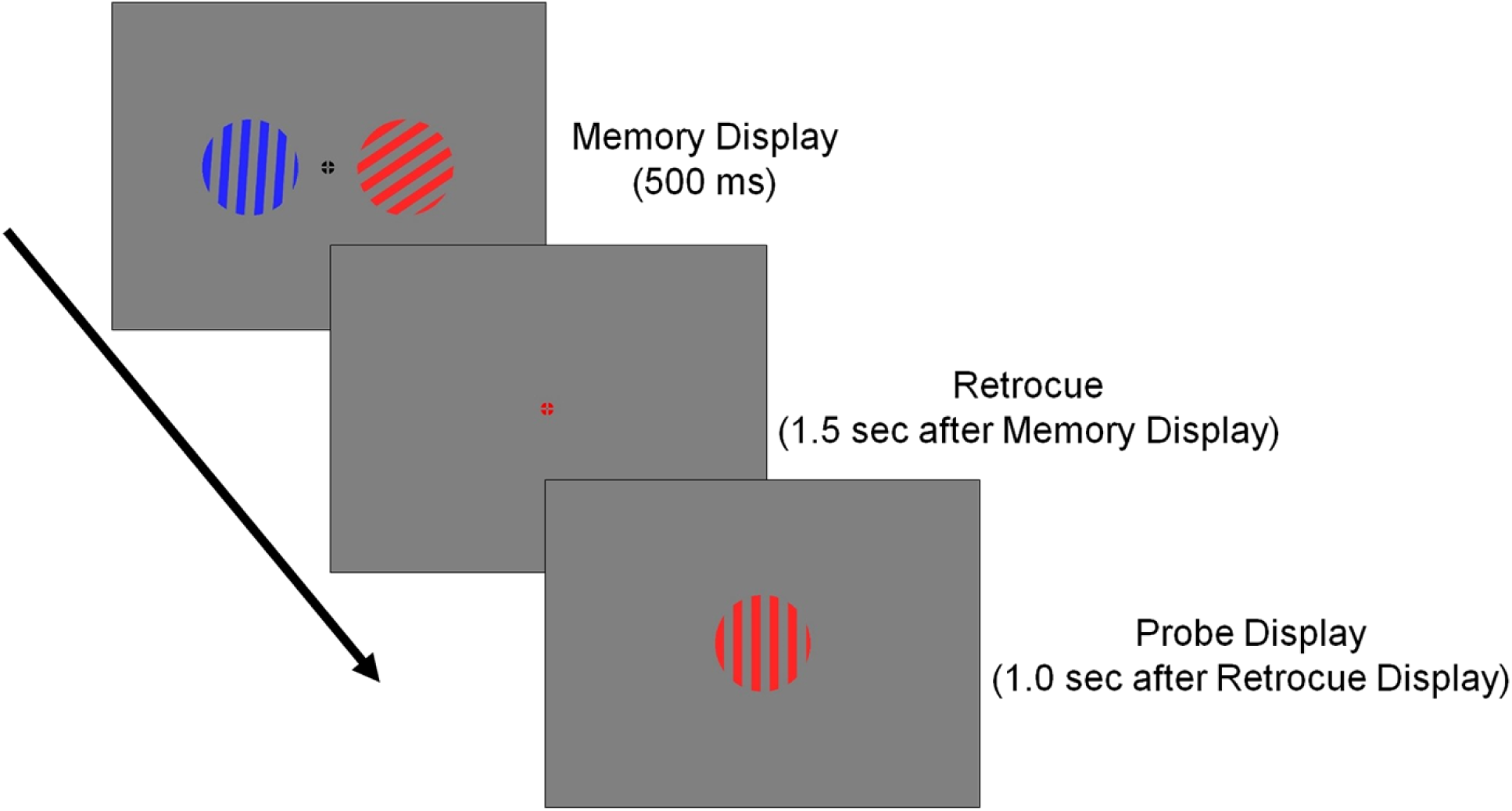
Experimental Task. Participants remembered the orientations of two gratings over a blank delay, then recalled the orientation of a retrospectively cued grating. See text for details.

## Methods

### Sample Size Justification

Prior to data collection, an a priori sample size of 32 human adult volunteers (both sexes) was selected. This choice was based on published empirical findings and effect sizes (e.g., van Ede et al., 2019a).

### Research Participants

32 volunteers from the University of Nevada, Reno participated in this experiment. Each participant completed a single 2.5-hour testing session in exchange for monetary remuneration ($15/h). All volunteers self-reported normal or corrected-to-normal visual acuity and gave both written and oral informed consent before enrolling in the study. All study procedures were approved by the local institutional review board. Data from one participant who completed the study were excluded from final analyses due to chance-level task performance (i.e., the participant used the correct response hand on only 52% of trials); thus, the findings reported here reflect the remaining 31 participants.

### Testing Environment

Participants were seated in a dimly lit room for the duration of testing. Visual displays were generated in MATLAB and rendered on a 27-inch LCD monitor (1920 x 1080 resolution) cycling at 240 Hz using Psychtoolbox-3 (Kleiner et al., 2007). Participants were seated approximately 80 cm from the display (head position was not constrained). Participants responded via buttons on a standard US computer keyboard.

### Visuomotor Recall Task

A task schematic is shown in Figure 1. Each trial began with a 500 ms sample display containing two colored gratings (radius 4.5 degrees visual angle [DVA] from a viewing distance of 80 cm; 1.0 cycles/DVA) presented 7.0 DVA to the left and right of a central fixation point. The orientation of each grating on each trial was randomly and independently sampled (with replacement) from the set [20, 40, 60, 80, 120, 140, 160°), and a small amount of angular jitter (±1-5°) was added to each orientation on each trial to discourage verbal coding. Stimulus orientations were counterbalanced across the entire experiment (though not necessarily within a single block of trials).

The sample display was followed by a 1500 ms blank display and a 1000 ms retrocue display where the fixation cross changed colors from black to either blue or red. The color of the retrospective cue instructed participants which grating would be probed at the end of the trial (i.e., blue or red) and which hand should be used to make a behavioral response (left or right). Cue color-response mappings (i.e., red cue/left hand vs. red cue/right hand) were counterbalanced across participants. Each trial ended with a 2000 ms probe display containing a vertically oriented grating; participants recalled the orientation of the cued grating by adjusting the probe grating to match the corresponding sample grating using the cue-specific response hand. If participants were satisfied with their response before the end of the 2000 ms, they were instructed to refrain from pressing any other buttons. Participants completed 3 (N = 1), 7 (N = 2), or 8 (N = 28) blocks of 72 trials.

### EEG Acquisition & Preprocessing

Continuous EEG was recorded from 63 scalp electrodes using a BrainProducts actiCHamp system. Online recordings were referenced to the left mastoid (10-20 site TP9) and digitized at 1 kHz. The following offline preprocessing steps were applied, in order: (1) resampling from 1 kHz to 500 Hz; (2) high-pass filtering (0.5 Hz using zero-phase forward- and reverse-finite impulse response filters as implemented by EEGLAB software extensions; Delorme & Makeig, 2004); (3) identification and reconstruction of noisy electrodes and epochs via artifact subspace reconstruction (implemented via EEGLAB; Chang et al., 2019), (4) re-referencing to the average response of all electrodes, (5) epoching from -1.0 to +5.0 sec relative to the start of each trial, (6) detection and removal of oculomotor and motor artifacts via independent components analysis and automated EEGLAB artifact detection functions, and (7) application of a surface Laplacian to remove low spatial frequency components from the signal (Perrin et al., 1989). Omitting or altering any of these steps (e.g., re-referencing to the algebraic mean of the left and right mastoids or omitting artifact subspace reconstruction) had no qualitative impact on any of the findings reported in the manuscript.

### Eyetracking Data Acquisition and Preprocessing

We obtained high-quality binocular eye position data for 18 of 32 participants enrolled in the study. Eye position data were acquired using an SR Research Eyelink 1000 Plus infrared eyetracker operating in remote (i.e., “head-free”) mode and digitized at 500 Hz. The eyetracker was calibrated 2-3 times per testing session using a standard nine-point grid included with the tracker hardware. Eye position data were filtered for blinks and horizontal eye movements > 2 DVA, then epoched from -1000 to +2000 ms relative to retrocue onset. No other preprocessing steps were applied.

### Spectral EEG Analyses

Spectral analyses focused on occipitoparietal and frontocentral 10-20 electrode site pairs O1/2, PO3/4, PO7/8, C1/C2, and C3/C4; analyses were restricted to a period spanning -500 to +2500 around retrocue onset. We extracted broadband spectral power from each electrode on each trial using a short-time Fourier transform (STFT) with a frequency range from 1-40 Hz (in 0.25 Hz steps) using a 200 ms sliding window and 2 ms step size. Power was computed by squaring the absolute value of the complex Fourier coefficients within each STFT window. To quantify changes in lateralized activity during visual selection, we sorted power estimates at each occipitoparietal electrode site by the location of the retrospectively cued stimulus, i.e., contralateral vs. ipsilateral hemifield, and expressed this difference as a normalized percentage:

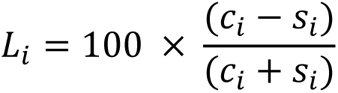

where c_i_ and s_i_ are the trial-averaged responses of electrode *i* when the retrospectively cued grating appeared in the contralateral or ipsilateral visual field, respectively. Lateralization estimates were pooled across occipitoparietal electrode sites, yielding a single time-frequency matrix of power estimates per participant. An identical approach was used to quantify changes in lateralized activity during motor selection, with the exceptions that (a) analyses focused exclusively frontocentral electrode sites, and (b) trial-wise power estimates were sorted by the retrospectively cued response hand rather than stimulus location.

To generate topographical maps of alpha- and beta-band lateralization, we repeated this analysis for every scalp electrode and averaged power estimates over 8-13 Hz (“alpha-band” activity) and 15-30 Hz (“beta-band” activity) over a period from 500-1000 Hz. These frequency bands were chosen a priori based on commonly reported values in the literature (e.g., Klimesch, 2012), and this specific temporal window was chosen a priori based on prior reports suggesting that it takes human observers 300-500 ms to process and respond to a retrospective memory cue (e.g., Souza et al., 2014).

To visualize changes in occipitoparietal alpha power, frontocentral mu-alpha power, and frontocentral beta power, we extracted and averaged lateralization estimates from 8-13 Hz over electrode sites O1/2, PO3/4, and PO7/8, lateralization estimates from 8-13 Hz over electrode sites C1/2, C3/4, and lateralization estimates from 15-30 Hz over electrode sites C1/2, C3/4. Statistical analyses of lateralization were performed using nonparametric sign permutation tests with temporal cluster-based correction (see *Statistical Analyses*, below).

### Quantifying Eye Position Biases

Preprocessed eye position data were sorted by the location of the retrospectively cued stimulus (i.e., left vs. right visual field). Since stimuli were rendered along the vertical meridian, we restricted our analyses to horizontal eye position measurements. Following earlier work (van Ede et al. 2019b), we computed a normalized measure of gaze bias by converting pixelwise recordings into a percentage along an axis extending from fixation to the center of each visual stimulus. Thus, a ±100% gaze bias would result from participants foveating the center of one stimulus, while a 0% gaze bias would result from participants perfectly holding fixation. Gaze biases during right visual field trials were sign reversed and averaged with biases from left visual field trials, yielding a single gaze position bias time course per participant.

### Orientation Decoding

Orientation decoding analyses were performed on broadband (0.5 Hz+) signals measured at occipitopartietal electrode sites O1/2, PO3/4, PO7/8 and frontocentral electrode sites C1/2, C3/4. Decoding performance was computed separately for each time sample using 10-fold cross-validation. During each cross-validation fold, we designated 90% of available trials as a training data set and the remaining 10% of trials as a test data set, taking care to ensure that the training data set contained an equal number of observations for each orientation. Decoding performance was estimated using the multivariate distance between EEG activity patterns associated with memory for specific experimental conditions (Wolff et al., 2017). We computed the Mahalanobis distance between trial-wise activation patterns in each test data set with position-specific activation patterns in the corresponding test data set, yielding a set of distance estimates. If scalp activation patterns contain information about stimulus orientation, then distance estimates should be smallest when comparing patterns associated with memory for similar or identical orientations in the training and test data sets and largest when comparing orthogonal orientations. Trial-wise distance functions were averaged and sign-reversed for interpretability. Decoding performance was estimated by convolving timepoint-wise distance functions with a cosine function, yielding a metric where chance decoding performance is equal to 0. Decoding results from each training- and test-data set pair were averaged (thus ensuring the internal reliability of our approach), yielding a single decoding estimate per participant, and timepoint. To facilitate interpretability, orientation decoding timeseries were smoothed with a 100 ms Gaussian kernel. We verified that smoothing did not qualitatively affect any of the findings reported in the manuscript.

### Statistical Comparisons

Statistical comparisons were based on nonparametric signed randomization tests (Maris & Oostenveld, 2007). Unless otherwise specified, each test we performed assumes a null statistic of 0 (i.e., no difference in alpha- or beta-band lateralization, or no difference in observed vs. chance decoding performance). We therefore generated null distributions by randomly relabeling each participant’s data with 50% probability and averaging the data across participants. This step was repeated 10,000 times, yielding a 10,000-element null distribution for each time point. Finally, we implemented a cluster-based permutation test with cluster-forming and cluster-size thresholds of p < 0.05 (two-tailed) to evaluate observed differences with respect to the null distribution while accounting for signal autocorrelation.

## Results

We recorded EEG while 31 human volunteers performed a retrospectively cued orientation memory task (Figure 1). Participants remembered the orientations of two gratings over a blank delay. 1.5 sec after encoding, a 100% valid color cue indicated which of the two gratings would be probed for report at the end of the trial and which hand should be used for recall (i.e., left or right; response hand/cue color mappings were counterbalanced across participants). A response display containing a vertically oriented grating was presented 1.0 seconds later; participants adjusted the orientation of the probe to match the orientation of the cued grating using the cued response hand. Participants performed this task well, recalling the orientation of the probed grating with an average (±1 S.E.M.) absolute error of 13.6° ± 0.75° and responding with the correct hand on 93.8% ± 0.89% of trials. The average response time (i.e., elapsed time between the onset of the probe display and the participant’s initial button press) was 451 ±14.9 msec.

### EEG Signatures of Spatial and Motor Selection

Following earlier work (van Ede et al., 2019a), we first identified EEG signals that co-varied with the physical location of the retrospectively cued stimulus (i.e., left vs. right visual hemifield) and EEG signals that co-varied with the retrospectively cued response hand (i.e., left vs. right). Because stimulus location and response hand were independently manipulated over trials (i.e., a stimulus requiring a right-hand response was equally likely to appear in the left vs. right visual field) we were able to characterize the selection of both attributes in trial-averaged EEG data independently of one another and any nuisance effects (e.g., volume conduction or other signal mixing). We therefore attributed patterns of neural activity that covaried with stimulus location to the selection of spatial information stored in WM and patterns of neural activity that covaried with response hand to the selection of motor information stored in WM.

Spatial selection was associated with a robust but transient decrease in 8-13 Hz alpha power over electrode sites contralateral to the location of the cued stimulus (Figure 2A). This modulation was strongest over occipitoparietal electrode sites (Figure 2A, inset), consistent with prior reports linking changes in lateralized alpha power to shifts of attention in perception and WM (Klimesch, 2012). Conversely, motor selection was associated with a robust and sustained decrease in 8-13 Hz mu-alpha power and 15-30 Hz mu-beta power over electrode sites contralateral to the cued response hand (Figure 2B). This modulation was strongest over frontocentral electrode sites (Figure 2B, inset), consistent with prior reports linking changes in lateralized mu-alpha and beta power to response preparation and execution. Direct comparisons of EEG signals associated with visual and motor selection revealed transient visual selection lasting approximately 250-850 ms and sustained motor selection lasting 250-3000+ ms after retrocue onset (Figure 2C). However, neither the onset, magnitude, nor duration of spatial or motor selection during the cue-to-probe interval varied as a function of participants’ response times (i.e., time to first keypress following probe onset; Figure 3A-B) nor participants’ memory performance (i.e., recall error; Figure 3C-D).

**Figure 2.**
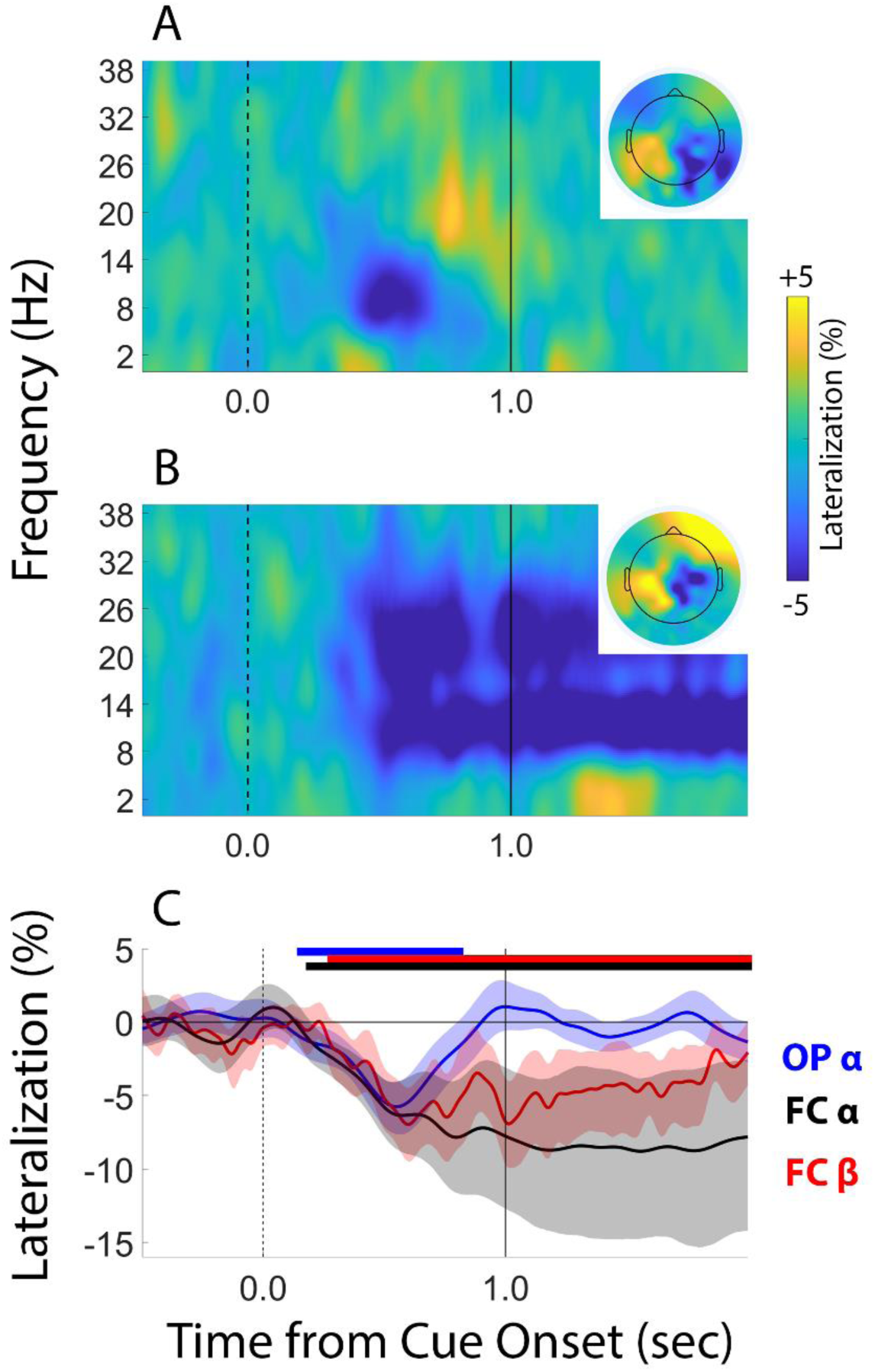
EEG Signatures of Spatial and Motor Selection. (A) Spectral signals that co-varied with stimulus location (i.e., left vs. right visual field). Lateralization estimates were computed from occipitoparietal electrode site pairs O1/2, PO3/4, and PO7/8. The inset shows the distribution of lateralized alpha power (8-13 Hz) across all scalp electrodes, averaged over a period spanning 0.5 to 1.0 sec after cue onset. (B) Spectral signals that co-varied with response demands (i.e., left vs. right hand). Lateralization estimates were computed from frontocentral site pairs C1/2 and C3/4. The inset shows the distribution of lateralized alpha/beta power (8-13 Hz) across all scalp electrodes, averaged over a period spanning 0.5 to 1.0 sec after cue onset. (C) Direct comparisons of occipitoparietal alpha (OP α), frontocentral mu-alpha (FC α) and frontocentral mu-beta (FC β) power. Horizontal bars at the top of the plot depict intervals where lateralization estimates were significantly less than 0%. Shaded regions depict the 95% confidence interval of the mean. Vertical bars at time 0.0 and 1.0 in each plot depict the onset of the retrocue and probe displays, respectively.

**Figure 3.**
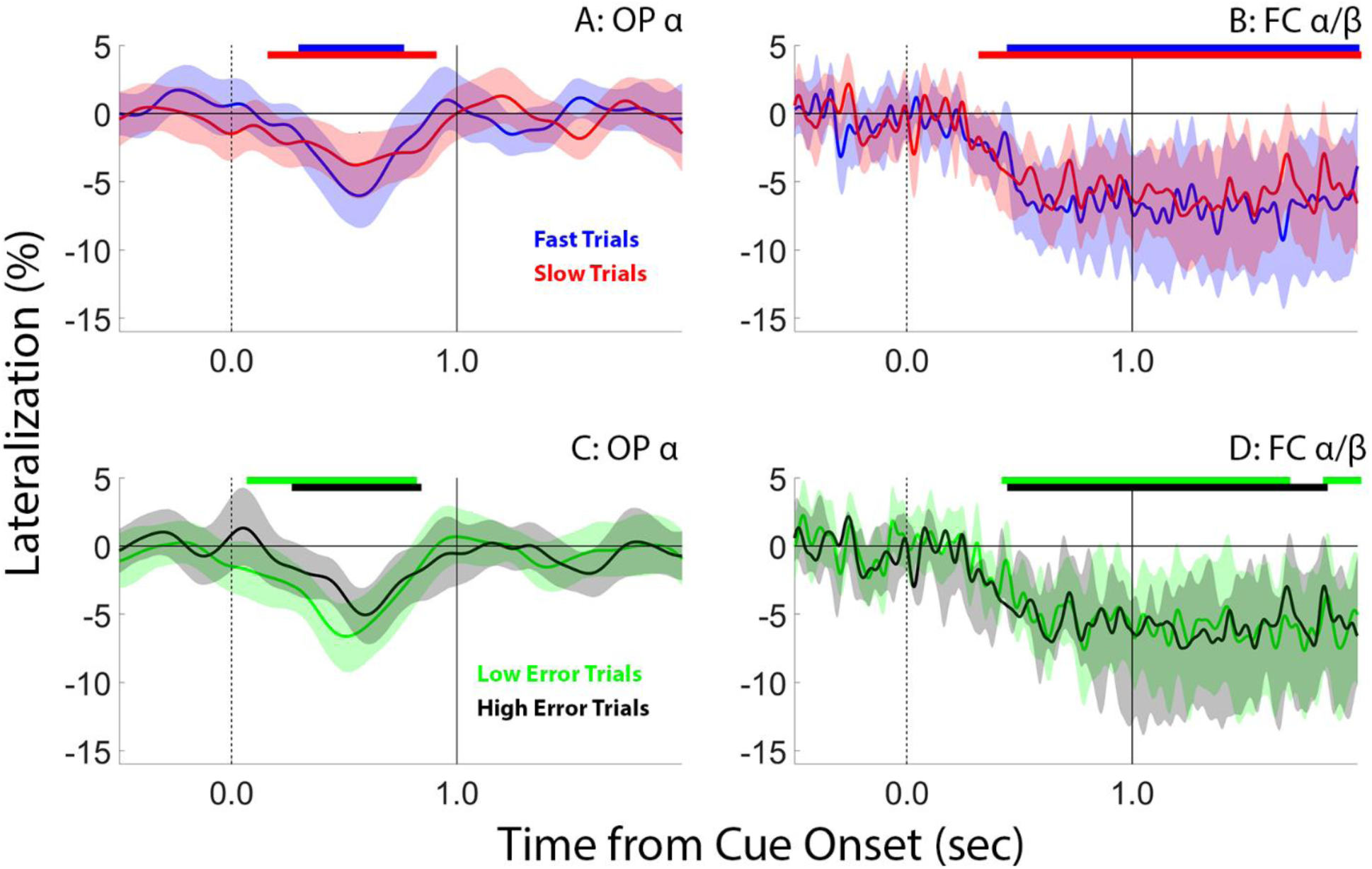
EEG signatures of spatial and motor selection do not predict trial-wise variability in task performance. Trial-wise differences in occipitoparietal (OP) α power (8-13 Hz; Panel A) and frontocentral (FC) α/β power (8-30 Hz) did not predict trial-wise differences in response onset (Panels A and B, respectively) or memory precision (Panels C and D, respectively). Horizontal bars at the top of each plot depict epochs where EEG lateralization was significantly less than zero (cluster-corrected permutation tests). Shaded regions depict the 95% confidence interval of the mean.

### Oculomotor Signatures of Spatial and Motor Selection

A handful of recent studies have reported small but robust biases in gaze (e.g., van Ede et al., 2019b) and head direction (Thom et al., 2023) towards the physical location of a retrospectively cued stimulus held in WM, even when location is task-irrelevant. Importantly, these gaze biases appear to be at least partially independent of EEG signals associated with covert spatial attention (e.g., Liu et al., 2022; see also Yu et al., 2022 for an analogous finding in nonhuman primates). Motivated by these findings, we sought to compare the amplitudes and latencies of EEG and oculomotor signals of covert spatial attention. We obtained high-quality horizontal gaze position recordings for 18 of the 31 participants who completed this study (EEG and gaze position data were acquired concurrently, see Methods). Following earlier work (van Ede et al. 2019b), we converted participants’ horizontal gaze position measurements into a normalized metric where 0% represents perfect fixation and -100% represents foveating the center of the cued stimulus (we used a negative scale to facilitate visual comparisons with negative-going changes in EEG lateralization; Figure 2). Consistent with earlier findings (e.g., van Ede et al. 2019b), we observed a small (∼3%) but robust bias in horizontal gaze position towards the cued stimulus beginning 282 ms after cue onset (Figure 4A).

**Figure 4.**
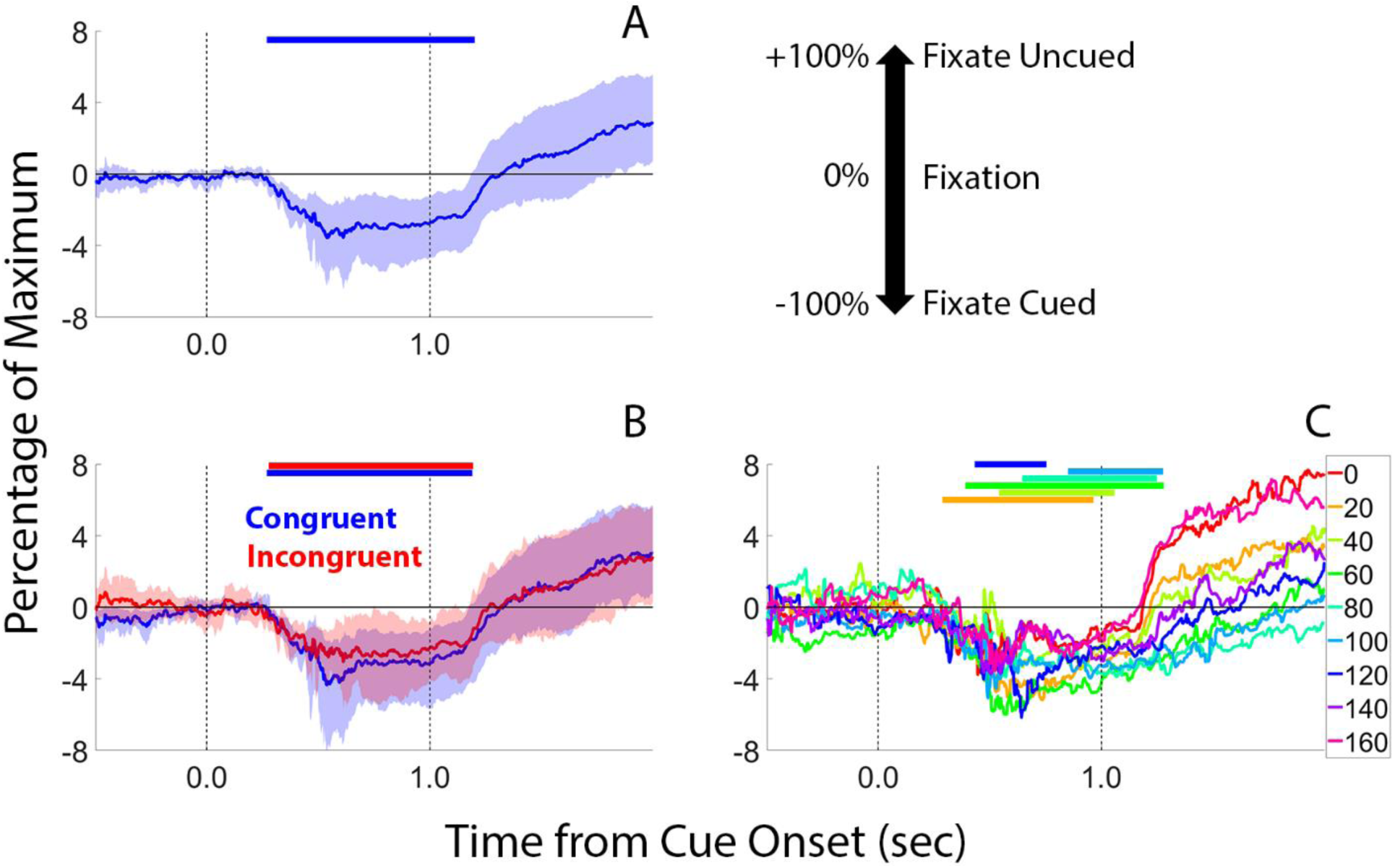
Horizontal Gaze Biases Associated with Spatial, Motor, and Feature Selection. (A) Even though the parameters of our task rendered spatial information irrelevant (i.e., participants could solve the task by remembering color-orientation bindings regardless of their positions during encoding), we observed a small but robust horizontal bias towards the position of the retrospectively cued stimulus. Gaze biases are expressed as a percentage of the maximum possible bias, with -100% corresponding to fixating the retrospectively cued grating and +100% corresponding to fixating the non-cued grating. Shifts towards the cued grating are plotted downwards to facilitate comparisons with the negative-going EEG lateralization estimates shown in Figure 2. (B) Gaze biases were equivalent in magnitude and duration during congruent trials where spatial and motor selection were aligned (e.g., a left visual field stimulus requiring a left-hand response) and incongruent trials where spatial and motor selection were misaligned (e.g., a left visual field stimulus requiring a right-hand response), suggesting that gaze biases were driven exclusively by the cued stimulus position. (C) Gaze biases during the cue-to-probe interval were unaffected by stimulus orientation. Shaded regions in Panels (A) and (B) depict the 95% confidence interval of the mean; shaded regions were omitted from Panel (C) to improve readability. Horizontal bars at the top of each plot depict intervals where gaze bias was significantly less than 0%. Vertical lines at times 0.0 and 1.0 depict the onset of the cue and probe displays, respectively.

In principle, gaze biases present during visuomotor selection could reflect the attentional selection of the cued stimulus’ position, the cued response hand, or some mixture of both. While our study was not designed to disentangle these possibilities, we reasoned that if gaze position biases are jointly influenced by spatial and motor selection then they should be larger during congruent trials where the selection of spatial and motor information is aligned (e.g., a left visual field stimulus requiring a left hand response) compared to incongruent trials where the selection of spatial and motor information is misaligned (e.g., a left visual field stimulus requiring a right hand response). However, direct comparisons of gaze biases during congruent and incongruent trials revealed no differences in either the magnitude nor the duration of horizontal gaze biases (Figure 4B), suggesting that these biases are driven primarily by the spatial position of the to-be-recalled stimulus and not the preparation of a manual response.

Finally, we considered the possibility that gaze position biases were jointly influenced by stimulus position and orientation. We tested this possibility by replotting and analyzing horizontal gaze bias as a function of stimulus position (left vs. right hemifield) and stimulus orientation (0°-160° in 20° increments; Figure 4C). This analysis revealed a significant modulation of gaze position biases by stimulus orientation during the later portion of the recall period (250-500 ms after probe onset; one-way repeated measures analysis-of-variance with stimulus orientation as the sole factor; F(8,136) = 3.995, p = 0.0003, η^2^ = 0.190), but not during the early portion of the recall period (0-250 ms after probe onset; F(8,136) = 1.412, p = 0.197, η^2^ = 0.077) nor during the cue-to-probe interval (e.g., 500-1000 ms after retrocue onset; F(8,136) = 0.778, p = 0.623, η^2^ = 0.044). Like the EEG data, gaze biases during the cue-to-probe interval did not predict participants’ response speeds (i.e., time to first keypress following probe onset; Figure 5A) nor participants’ memory performance (i.e., recall error; Figure 5B).

**Figure 5.**
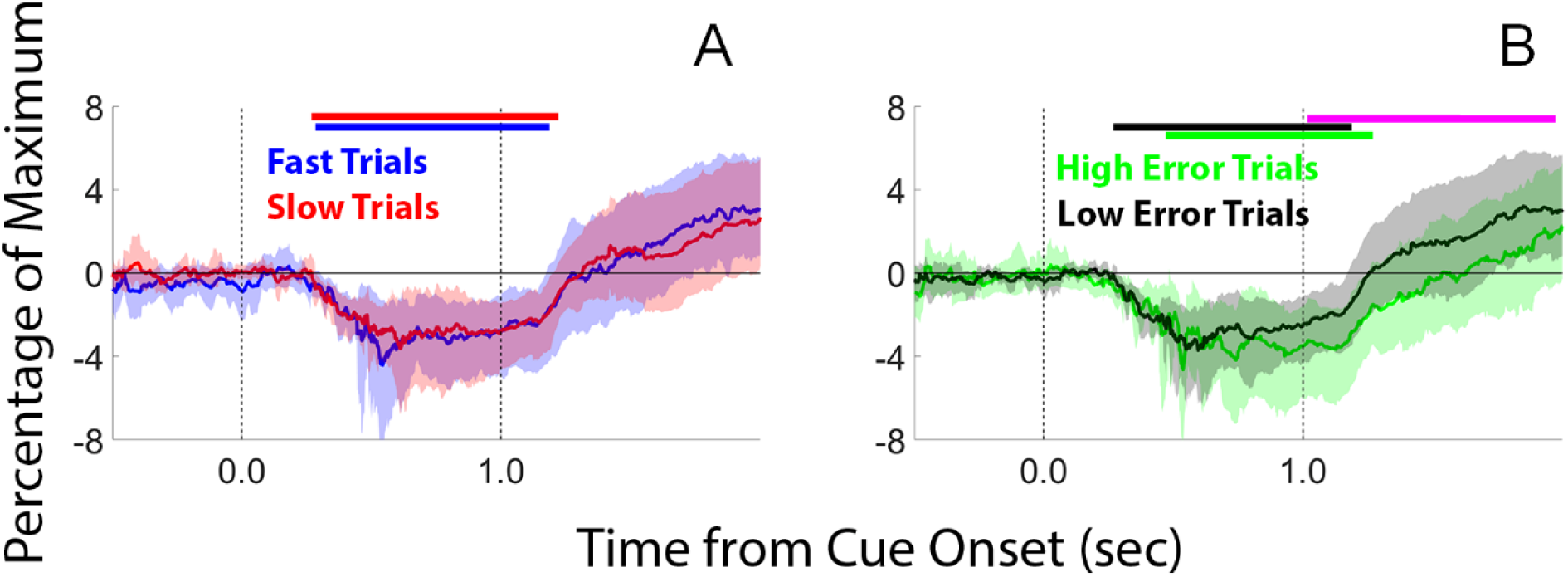
Oculomotor signatures of spatial selection during the cue-to-probe interval do not predict trial-wise variability in task performance. Trial-wise differences in gaze bias did not predict variability in response onset (A) during either the cue-to-probe or recall periods. Trial-wise differences in gaze bias did not predict variability in recall error (B) during the cue-to-probe interval but did predict recall error during the recall period, with low-error trials associated with reliably smaller gaze position biases during the recall period compared to high-error trials. Horizontal bars at the top of each plot depict epochs where gaze position was significantly less than zero (i.e., towards the location of the retrospectively cued disc; cluster-corrected permutation tests). Shaded regions depict the 95% confidence interval of the mean.

### EEG Signatures of Feature Selection

Several lines of evidence indicate that the brain automatically retains spatial information in WM: First, location-specific representations can be reconstructed from scalp EEG activity during WM tasks where location is irrelevant (Foster et al., 2017). Second, human gaze (e.g., van Ede et al., 2019b) and head position (Thom et al., 2023) are biased towards the location of a behaviorally prioritized item stored in WM, also in tasks where location is irrelevant. Third, changes in gaze position or shifts of covert spatial attention during WM can disrupt or introduce systematic biases in memory for visual features (e.g., color; Golomb et al., 2014), and memory guided comparisons for successively presented visual stimuli are impaired when they are rendered in different spatial positions (e.g., Hollingworth, 2007).

Automatic storage of visuospatial information in WM may be adaptive, enabling the segregation of mental representations in retinotopic or spatiotopic coordinate frames during memory encoding, storage, and retrieval (Groen et al., 2022). Our data support this perspective: even though our experimental task (Figure 1) did not oblige participants to retain spatial information in WM, we nevertheless observed robust electrophysiological (Figure 2) and oculomotor (Figure 4) evidence for the selection of spatial information following the appearance of a retrospective cue.

We reasoned that if visuospatial information plays an anchoring function during WM storage and WM-guided behavior, then the selection of spatial information might also prompt the selection of task-relevant feature information, in this case, orientation. We examined this possibility by testing whether it was possible to decode the orientations of the cue-matching and -nonmatching gratings during the cue-to-probe interval when evidence for spatial selection was highest (e.g., Figure 2C). Surprisingly, although we observed clear EEG and oculomotor evidence for spatial (and motor) selection during the cue-to-probe interval, above-chance decoding of the retrospectively cued orientation only emerged during the recall period (Figure 6). This general pattern held when we decoded stimulus orientation from the same occipitoparietal electrodes used to quantify spatial selection (O1/2, PO3/4, and PO7/8; Figure 2C= and Figure 6A), the same frontocentral electrode sites used to quantify motor selection (FC1/2; FC3/4; Figure 2C and Figure 6B), or all 62 scalp electrodes (Figure 6C).

**Figure 6.**
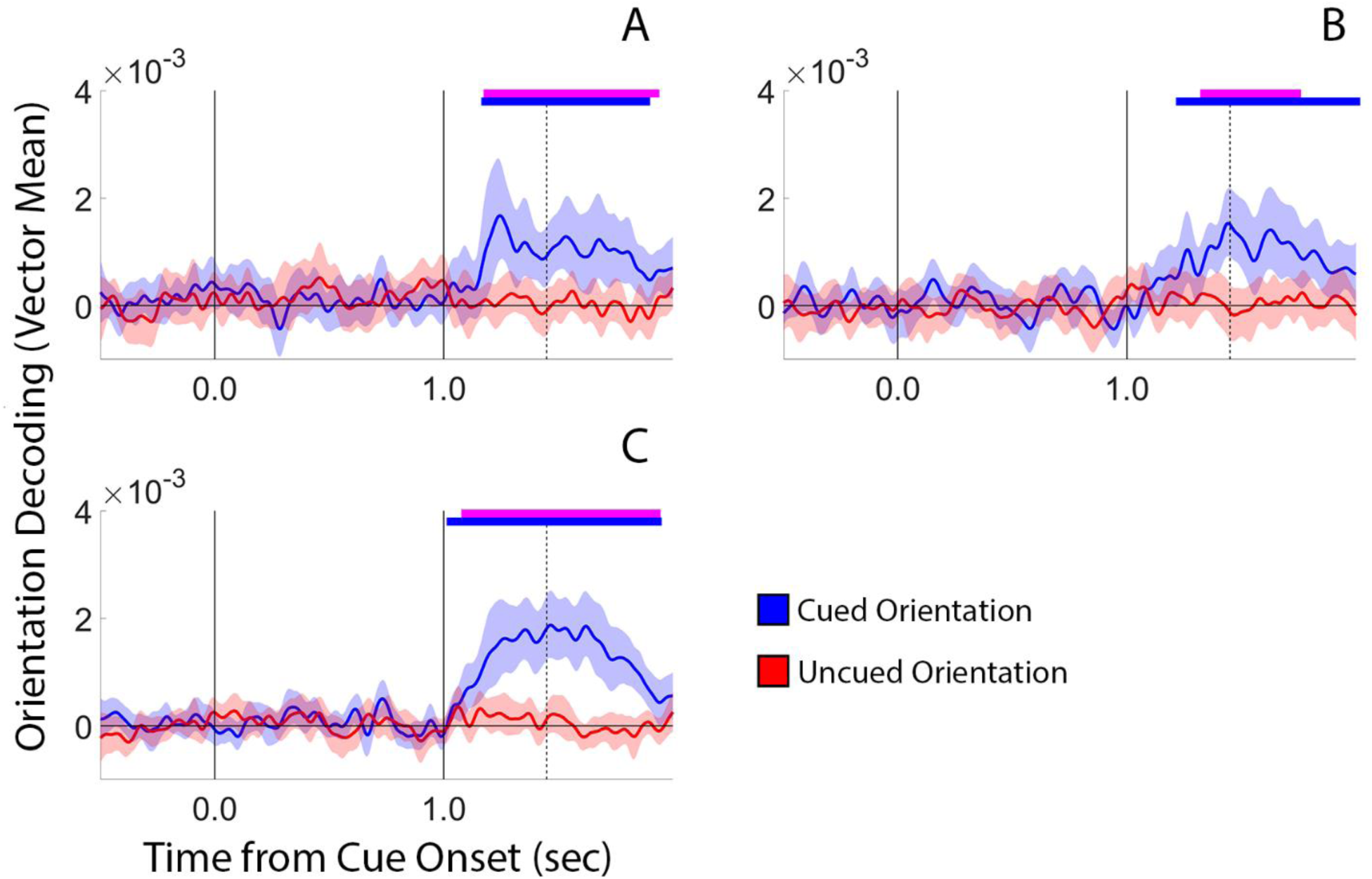
Orientation Decoding Time-Locked to Retrocue Onset. Decoding was computed from occipitoparietal electrode site pairs O1/2, PO3/4, and PO7/8 (A), frontocentral site pairs FC1/2 and FC3/4 (B), or all 62 scalp electrodes that we recorded from (C). Horizontal bars at the top of each plot mark epochs where decoding performance was significantly greater than chance; pink horizontal bars mark epochs where cued orientation decoding performance was significantly greater than uncued orientation decoding performance. The dashed vertical line at time 1.451 sec marks participants’ average response onset, i.e., the point at which they first began to rotate the probe grating. Shaded regions in each plot depict the 95% confidence interval of the mean.

We considered several explanations for the absence of above-chance orientation decoding during the cue-to-probe interval, as well as the presence of above-chance orientation decoding during the recall interval. First, we considered the possibility that above-chance decoding of orientation during the recall period reflects participants’ adjusting the orientation of the probe grating to match the orientation of the cued grating rather than the selection of orientation information stored in WM per se. We tested this possibility by comparing the onset of above-chance orientation decoding performance with the timing of participants’ responses (i.e., the amount of time that elapsed between the onset of the probe display and the time at which participants began to adjust the probe stimulus, i.e., 451 ±14.9 ms). The onset of above-chance decoding performance occurred significantly earlier than the average response onset for the data shown in Figure 6A (mean above-chance decoding onset relative to the appearance of the probe display = 182 ms; p = 0.0060; bootstrap test against a distribution with a mean of 451 ms), Figure 6B (mean above-chance decoding onset = 234 ms; p = 0.0004), and Figure 6C (mean above-chance decoding onset = 30 ms; p = 0.0001). Thus, above-chance decoding of orientation cannot be explained by participants adjusting the probe grating to match that of the cued grating.

Second, we considered the possibility that above-chance orientation decoding during the early portion of the response period (i.e., prior to response onset) was driven by the mere appearance of an oriented stimulus. However, note that the probe grating was always rendered with a vertical orientation, and thus, across trials the initial orientation of the probe grating was randomized with respect to the orientation of the remembered grating [drawn from the set [0°, 20°, 40°, 60°, 80°, 100°, 120°, 140°, 160°)]. This argues against the possibility that above-chance decoding during the initial part of the recall period was driven by the mere appearance of an oriented stimulus (we return to this point in the discussion).

Third, we considered the possibility that the absence of above-chance orientation decoding during the cue-to-probe interval – when participants were free to select the cue-matching orientation for upcoming report and when EEG and oculomotor evidence for spatial and motor selection was evident (e.g., Figures 2 and 4) – was due to a lack of analytic sensitivity. For example, one recent study reported that the orientation of a remembered stimulus could be decoded from alpha-band but not broadband EEG activity during WM storage (Barbosa et al., 2021). We therefore repeated our decoding analysis after applying an 8-13 Hz bandpass filter to the EEG data. The results of this analysis were remarkably similar to those summarized in Figure 6, with robust above-chance decoding of orientation observed during the recall period but not the cue-to-probe period (Figure 7).

**Figure 7.**
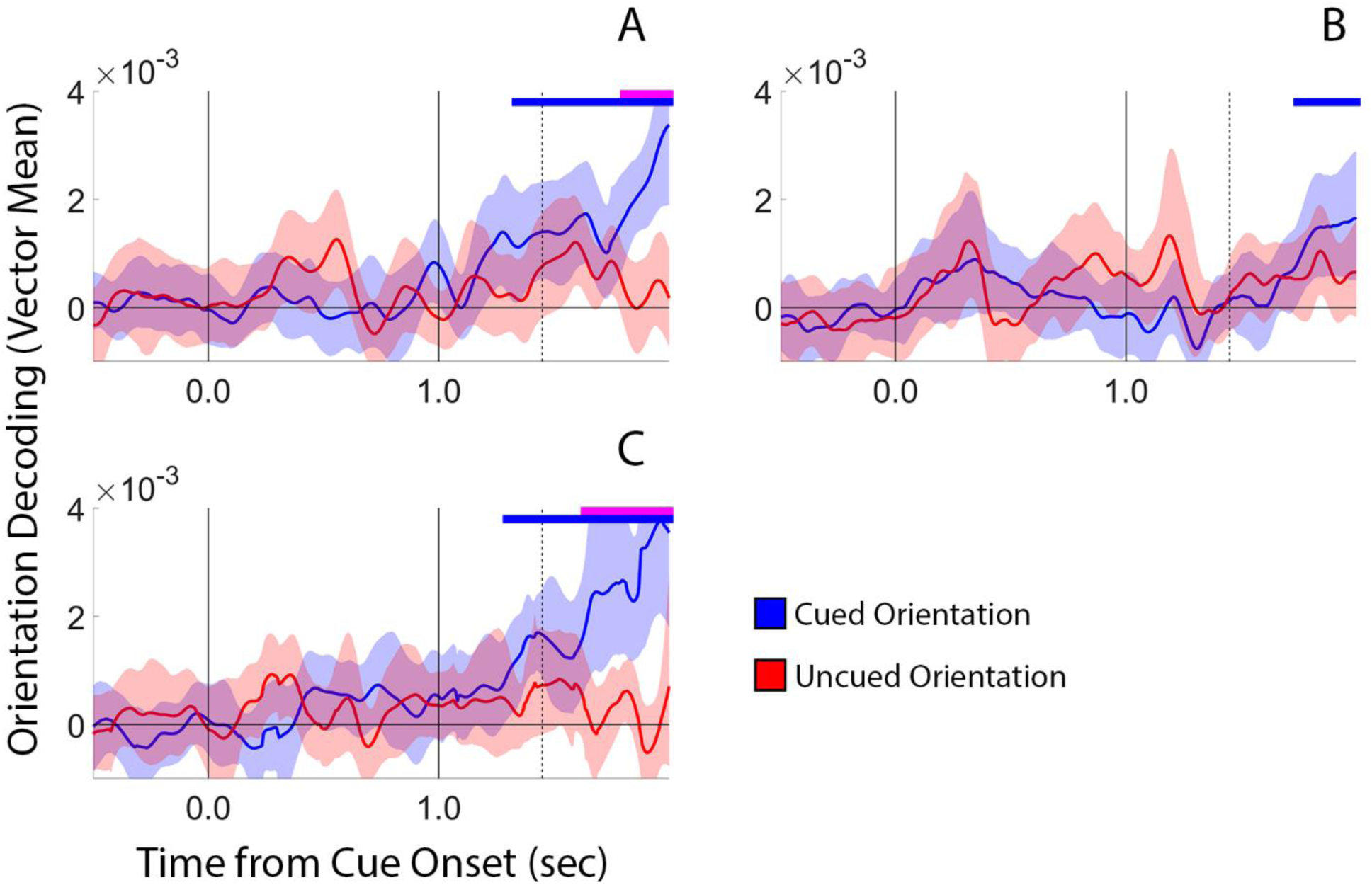
Orientation Decoding Performance Computed from Alpha-band EEG Signals (8-13 Hz). Conventions are identical to Figure 6.

Finally, we considered the possibility that the absence of above-chance orientation decoding during the cue-to-probe interval was caused by an analytic artifact. For example, one recent study demonstrated that high-pass filter cutoffs over 0.1 Hz can systematically bias the time(s) at which feature information can be decoded from broadband EEG data (van Driel et al., 2021). We tested this possibility by re-preprocessing our data with a 0.1 Hz high-pass filter cutoff and applying our decoding analysis to the resulting data. The results of this analysis were remarkably similar to the data summarized in Figures 6-7 (Figure 8), suggesting that the absence of above-chance decoding performance during the cue-to-probe interval cannot be explained by a filtering artifact.

**Figure 8.**
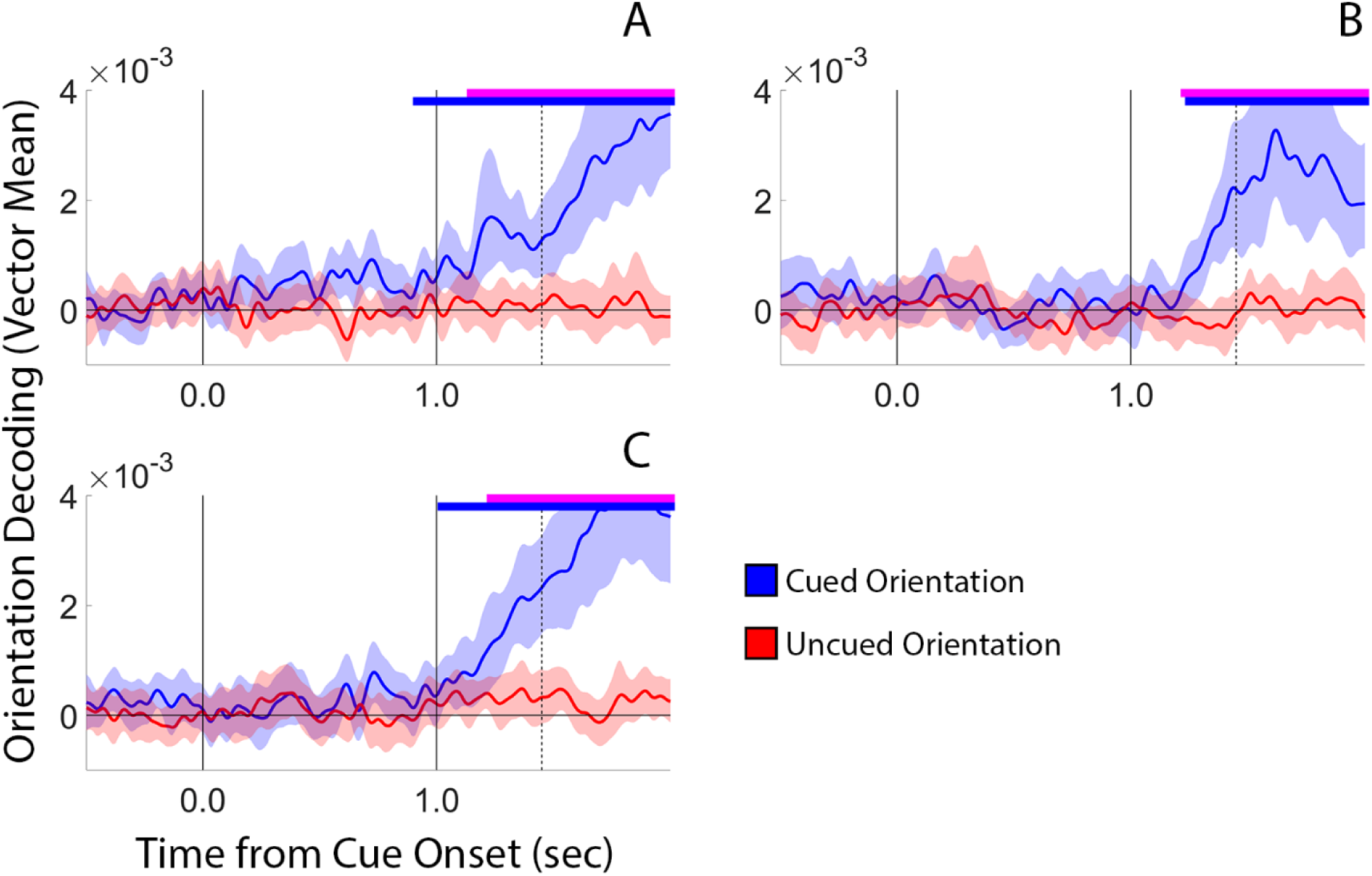
Orientation decoding time-locked to retrocue onset in 0.1 Hz high-pass filtered broadband EEG data. Conventions are identical to Figure 6.

### Eye Movement Control Analyses

Recent studies suggest that subtle differences in gaze position can contribute to feature decoding performance during WM storage and readout (e.g., Mostert et al., 2017; Quax et al., 2018). We therefore sought to understand whether and how gaze position may have influenced EEG decoding in the current study. As a first approximation, we tested whether horizontal gaze position biases were modulated by stimulus orientation during the cue-to-probe interval (i.e., -250 to -1 ms prior to probe onset) and during the recall period (i.e., 1 to 250 ms after probe onset). We averaged participants’ horizontal gaze position estimates as a function of stimulus orientation and subjected the resulting values to a one-way repeated-measures analysis of variance (ANOVA) with stimulus orientation as the sole factor. This analysis did not reveal an effect of orientation during the cue-to-probe interval (when it was not possible to decode stimulus orientation from EEG signals; F(8,136) = 0.778, p = 0.623, η^2^ = 0.044) or the recall period (when it was possible to decode stimulus orientation from EEG signals; F(8,136) = 1.412, p = 0.197, η^2^ = 0.077). However, additional analyses did reveal a robust effect of stimulus orientation on horizontal gaze position during later portions of the recall period (e.g., 250-500 ms after probe onset; F(8,136) = 3.995, p = 0.0003, η^2^ = 0.190). Thus, we undertook additional analyses to understand how differences in gaze position may have influenced EEG decoding performance.

To obtain a more direct estimate of the relationship between gaze position and stimulus orientation, we attempted to decode the latter from the former. Specifically, we used the same parametric distance-based decoding procedure used to decode stimulus orientation from EEG signals to decode stimulus orientation from trial- and time-wise records of gaze position in [x, y] screen coordinates. Due to a hardware malfunction vertical gaze position data from one participant were lost during recording. Thus, we restricted our analysis to the remaining 17 participants with concurrent and robust EEG and [x, y] gaze position recordings. This analysis revealed significant above-chance gaze position-based decoding of cued orientation during the recall period (Figure 9).

**Figure 9.**
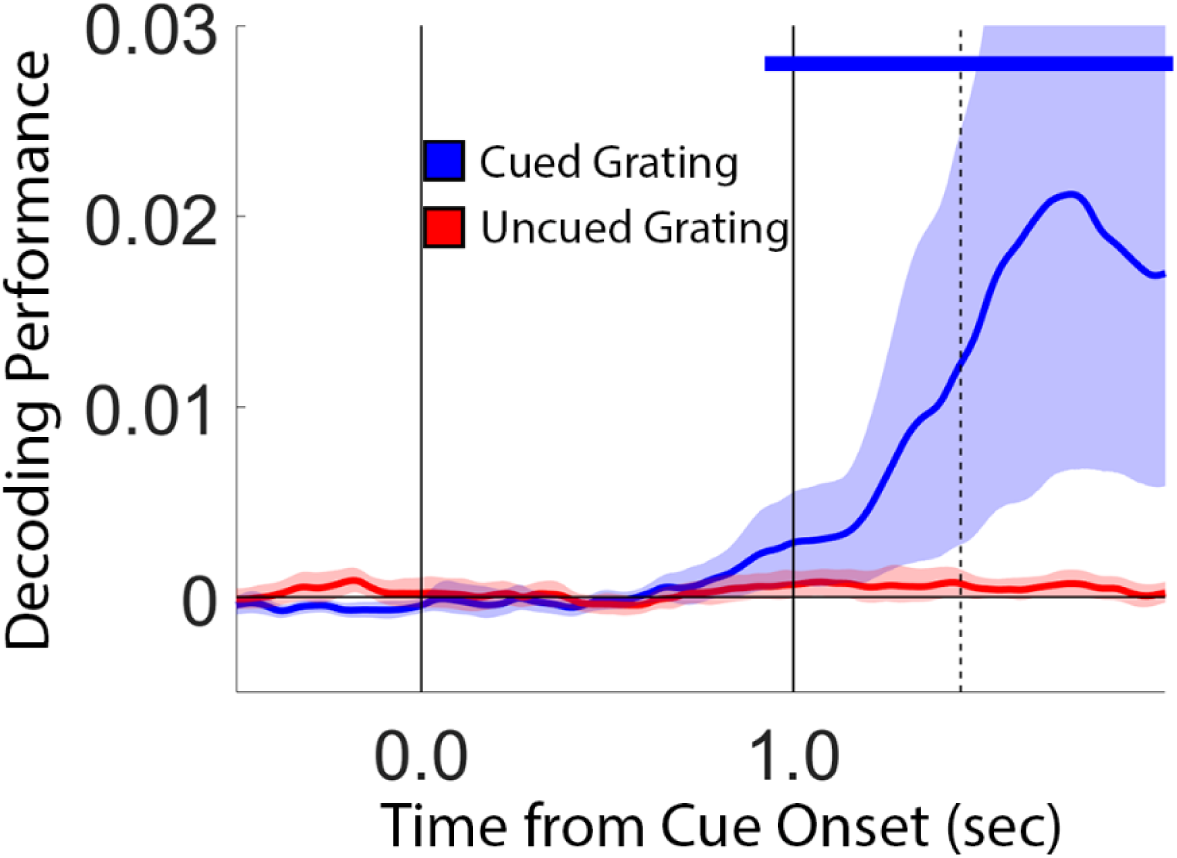
Orientation Decoding Performance Computed Using [X,Y] Gaze Position Data. Vertical lines at times 0.0 and 1.0 depict sec. the onset of the retrocue and probe displays, respectively. Vertical dashed line at time 1.451 sec. depicts the average response onset across participants. The blue horizontal bar at the top of the plot marks epochs where decoding performance was significantly greater than chance. Shaded regions around each line depict the 95% confidence interval of the mean. See text for additional details.

However, an examination of individual subject data indicated that above-chance decoding driven primarily by 3 extreme participants (Figure 10A, red traces). Following earlier work (Printzlau et al., 2022), we examined the influence of these participants on gaze position-based orientation decoding performance by computing average decoding performance during the probe-to-response period (i.e., 0 to 451 ms after probe onset, during which time it was possible to decode stimulus orientation from EEG data; Figure 6) while including and excluding suspected outliers from the analysis. As expected, gaze position-based orientation decoding performance fell to chance levels when the three extreme participants highlighted in Figure 10A were excluded from the analysis (Figure 10B; two-tailed t-test against chance, i.e., 0: t(13) = 1.96, p = 0.072). Finally, we confirmed that EEG-based decoding performance during the probe-to-response interval remained well above chance levels even when participants with extremely high gaze position-based decoding performance were excluded from the analysis (Figure 10C), indicating that gaze position biases were not solely responsible for above-chance EEG-based orientation decoding during this period.

**Figure 10.**
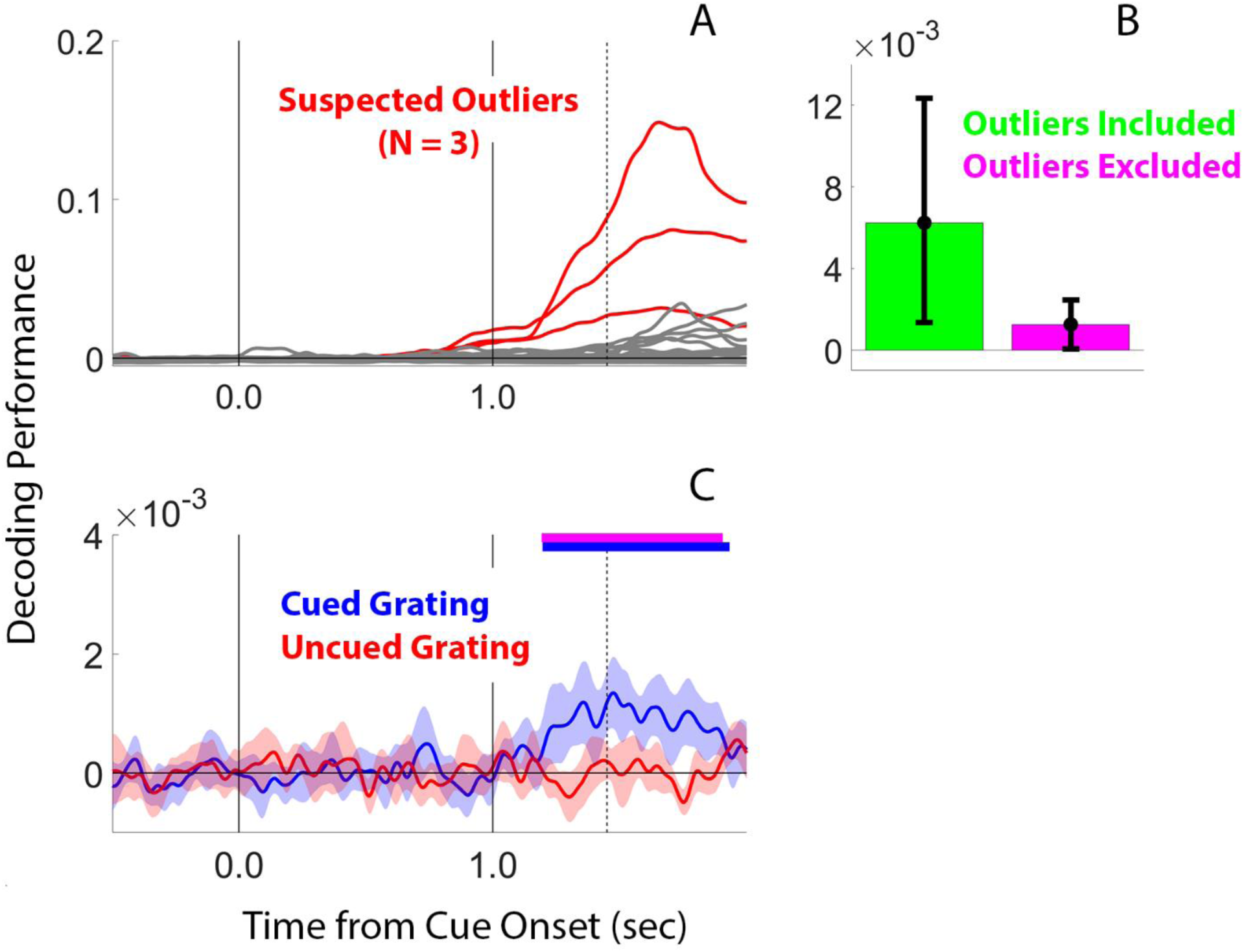
Influences of Gaze Position on Orientation Decoding from EEG Data. (A) A participant-by-participant analysis of orientation decoding computed from gaze position data (i.e., Figure 9) indicated that above-chance decoding performance was driven primarily by three extreme participants, highlighted here in red. (B) Removing the extreme participants shown in panel A reduced gaze position decoding of stimulus orientation during the probe-to-average response interval (i.e., 0 to 451 ms after probe onset) to chance levels. (C) To test whether gaze position biases were responsible for above-chance decoding of orientation from EEG data, we replotted EEG-based decoding performance from the 14/17 participants with concurrent EEG and eye tracking records (i.e., after excluding the three outliers highlighted in panel A) using all 62 scalp electrodes (e.g., compare with Figure 6C). It remained possible to decode the orientation of the cued grating from EEG data during the probe-to-response interval even though orientation decoding computed from gaze position data during the same interval was at chance. Vertical lines at times 0.0 and 1.0 depict sec. the onset of the retrocue and probe displays, respectively. Vertical dashed line at time 1.451 sec. depicts the average response onset across participants. The blue horizontal bar at the top of the plot marks epochs where decoding performance was significantly greater than chance. Shaded regions around each line depict the 95% confidence interval of the mean.

## Discussion

Human memory systems exist so that prior experiences can guide immediate and future behaviors. In recognition of this fact, recent conceptualizations of short-term and/or working memory have begun to emphasize the action-oriented nature of this system. Strikingly, recent experimental findings suggest that the human brain can store multiple stimulus-response mappings in WM, and that behaviorally relevant WM contents can be selected in parallel with required actions (van Ede et al., 2019a). Although this result seems to challenge classic “think-then-act” conceptualizations of human memory performance (e.g., Donders 1868/1969; Sternberg 1969), in our view the evidence supporting this challenge is limited. Here, we re-examined the chronometry of spatial, feature, and motor selection during WM-guided behavior during a task optimized to disentangle neural signals associated with these processes. We found clear evidence for a temporal dissociation between feature and visuomotor selection: while EEG signatures linked to spatial and motor selection appeared coincidentally shortly after the appearance of an informative recall cue (Figure 2), EEG signatures linked to feature selection appeared significantly later (Figure 6). Our findings reveal an important distinction between spatial and non-spatial selection during WM-guided behavior and clarify how these selection mechanisms align with the selection of item-specific motor plans. More generally, our findings challenge conceptualizations of WM-guided behavior based exclusively on serial vs. parallel visuomotor processing.

Intuitively, the visual features of a task-relevant object could be selected automatically along with its location, complementing demonstrations of object-based selection in WM where selecting one non-spatial attribute of an object (e.g., color) leads to the selection of other non-spatial attribute of the same object (e.g., orientation; Printzlau et al., 2022). Alternatively, given the apparent primacy of spatial position in organizing visual perception and WM (e.g., Groen et al., 2022), the selection of feature information stored in WM might be functionally and/or temporally decoupled from the selection of spatial information stored in WM. Our data support this view. Specifically, although we observed clear EEG and eye position evidence for spatial selection during the cue-to-probe interval (Figures 2 and 4), evidence for feature selection was limited to the recall period (Figure 6). Thus, our data reveal the existence of temporally distinct mechanisms for the selection of spatial and non-spatial visual information during WM-guided behavior.

In the current study, robust above-chance decoding of orientation was only possible during the recall period (Figure 6). One possibility is that the read-out of orientation information from WM can only proceed after the appearance of a trailing sensory stimulus. Activity-silent models of WM propose that information is stored via short-term changes in synaptic weights in a way that permits trailing sensory inputs to reactivate neural firing patterns seen during WM encoding (e.g., Mongillo et al., 2008; Stokes, 2015; Wolff et al., 2017). It is unclear, however, whether short-term changes in synaptic weights are restricted to the retinotopic locations where remembered stimuli appeared. In prior studies supporting activity-silent WM, trailing stimuli used to reactivate latent WM representations always appeared in the same retinotopic locations as remembered stimuli (e.g., Wolff et al. 2017); in the current study the remembered stimulus and the probe stimulus were always rendered in different retinotopic locations (i.e., laterally vs. foveally; respectively; Figure 1). One way to reconcile this apparent discrepancy is to propose that activity-silent WM is spatially global, i.e., the changes in synaptic weights responsible for storing information extend beyond the retinotopic locations where remembered stimuli appeared. This arrangement would dovetail with other observations of spatially global WM, where feature information can be decoded from neural populations retinotopically mapped vs. not retinotopically mapped to the location of a remembered stimulus (e.g., Ester et al., 2009).

Individual variability in WM is strongly correlated with general cognitive function (e.g., IQ; Cowan, 2001), and studies have shown that participants with high WM function efficiently gate access to this system (e.g., Feldman-Wustefeld & Vogel, 2019). Given that WM is a fundamentally action-oriented system, the ability to efficiently select content already in WM and use this information for behavior is likely to play an equally important role in cognitive performance. Here, we used EEG and eye position data to track the selection of spatial, feature, and motor information during WM-guided behavior. Our data reveal the presence of multiple temporally dissociable selection mechanisms for WM-guided behavior, for example, with spatial and motor selection occurring prior to the read-out of feature information from WM. More generally, our findings refute conceptualizations of WM-guided behavior based purely on serial vs. parallel visuomotor processing.

